# Sex Differences in Human Brain Structure at Birth

**DOI:** 10.1101/2024.06.20.599872

**Authors:** Yumnah T. Khan, Alex Tsompanidis, Marcin A. Radecki, Lena Dorfschmidt, APEX Consortium, Topun Austin, John Suckling, Carrie Allison, Meng-Chuan Lai, Richard A. I. Bethlehem, Simon Baron-Cohen

**Affiliations:** Department of Psychiatry, University of Cambridge, Cambridge, CB2 8AH; Autism Research Centre, Department of Psychiatry, University of Cambridge, Cambridge, UK, CB2 8AH; Social and Affective Neuroscience Group, IMT School for Advanced Studies Lucca, Lucca, Italy; Department of Psychiatry, University of Pennsylvania, Philadelphia, PA 19104, USA; Lifespan Brain Institute, The Children’s Hospital of Philadelphia and Penn Medicine, Philadelphia PA 19139, USA; Neonatal Intensive Care Unit, Cambridge University Hospitals NHS Foundation Trust, Cambridge, UK, CB2 0QQ; Peterborough Foundation NHS Trust, Cambridge, CB2 8SZ, UK; Campbell Family Mental Health Research Institute, Centre for Addiction and Mental Health, Toronto, Ontario, Canada; Department of Psychiatry, The Hospital for Sick Children, Toronto, Ontario, Canada; Department of Psychiatry, Temerty Faculty of Medicine, University of Toronto, Toronto, Ontario, Canada; Department of Psychology, Faculty of Arts and Science, University of Toronto, Toronto, Ontario, Canada; Department of Psychiatry, National Taiwan University Hospital and College of Medicine, Taipei, Taiwan; Department of Psychology, University of Cambridge, Cambridge, CB2 3EB, UK

## Abstract

Sex differences in human brain anatomy have been well-documented; however, their underlying causes remain controversial. Neonatal research offers a pivotal opportunity to address this long-standing debate. Given that postnatal environmental influences (e.g., gender socialisation) are minimal at birth, any sex differences observed at this stage can be more readily attributed to prenatal influences. Here, we assessed on-average sex differences in global and regional brain volumes in 514 newborns (236 birth-assigned females and 278 birth-assigned males) using data from the developing Human Connectome Project. On average, males had significantly larger intracranial and total brain volumes, even after controlling for birth weight. After controlling for total brain volume, females showed higher total cortical gray matter volumes, whilst males showed higher total white matter volumes. After controlling for total brain volume in regional comparisons, females had increased white matter volumes in the corpus callosum and increased gray matter volumes in the bilateral parahippocampal gyri (posterior parts), left anterior cingulate gyrus, bilateral parietal lobes, and left caudate nucleus. Males had increased gray matter volumes in the right medial and inferior temporal gyrus (posterior part) and right subthalamic nucleus. Effect sizes ranged from small for regional comparisons to large for global comparisons. While postnatal experiences likely amplify sex differences in the brain, our findings demonstrate that several global and regional on-average sex differences are already present at birth.

## Introduction

Note on terminology: Throughout this paper, all references to “sex differences” are intended to reflect differences observed in group averages.

While sex differences in human brain anatomy are well-evidenced (for a meta-analysis, see 1), their magnitude, significance, and implications remain a matter of substantial ongoing debate (for recent discussions, see 2, 3). Most notably, their underlying causes are a central point of scientific discussion and remain poorly understood. This area of research is of high importance because the prevalence of various psychiatric, neurological, and neurodevelopmental conditions differs by biological sex (4, 5). Given that variations in brain development are implicated in these conditions and overlap with neurobiological sex differences, it is likely that sex differences play a key role in the development of these conditions (6, 7, 4). A better understanding of sex differences, their underlying causes, and their onset could therefore help tailor diagnostic, prognostic, and support strategies to facilitate optimal health outcomes.

Sex differences in brain structure are hypothesised to arise from a complex interplay between biological and environmental factors regulating brain development (8). For instance, during the first and second trimesters of pregnancy, male fetuses produce around 2.5 times more testosterone than females fetuses (9). This prenatal surge in testosterone is understood as a key early biological mechanism instigating the sexual differentiation of the body and brain (10). Gender socialisation also begins early in childhood, leading to divergent life experiences for males and females that likely influence the lifespan development of the brain. As it is now recognised that both nature and nurture play important roles, the question is not whether sex differences are due to biology *or* environment, but rather the extent and ways through which each factor contributes to a given sex difference in the brain. While disentangling the relative contributions of these factors is a considerable challenge, studying sex differences at birth offers a partial but important solution. At birth, postnatal environmental influences such as gender socialisation are relatively minimal. Thus, if a given sex difference is present at birth, it can be more readily attributed to prenatal factors.

The neonatal period is typically defined as the first 4 weeks of life, and existing studies in the field typically involve infants with mean post-birth ages that extend beyond the neonatal period (e.g., 33 days post-birth in 11). As a result, an understanding of sex differences immediately after birth remains extremely limited. The majority of existing research has shown that male infants have larger intracranial and total brain volumes than female infants (11-15), often even after accounting for birth weight. However, one study has reported no differences in intracranial or total brain volumes in 2-5-week-olds (16), contradicting these prior studies. Male infants are also reported to have larger total gray and white matter volumes, though these differences do not persist after accounting for the sex difference in intracranial volume (12, 13). However, when controlling for total brain volume rather than intracranial volume, another study has reported that 1-month-old males had larger total white matter volumes, whilst females had larger total gray matter volumes (15). These observed discrepancies emphasise the need for further research to clarify sex differences in the neonatal brain.

Research into brain regional sex differences is even more limited and inconsistent, complicating the identification of regions that show reliable sex differences during early development. When using region-of-interest volumetry, one study reported no regional sex differences in early infancy after controlling for intracranial volume (12). However, when using voxel-based approaches such as tensor- and deformation-based morphometry, other studies have reported various regional sex differences even after controlling for brain size (11, 12). For instance, male infants had increased gray matter volumes in the insula, middle temporal gyrus, fusiform gyrus, and hippocampus, whilst female infants had increased volumes in the dorsolateral prefrontal, motor, and visual cortices (12).

In summary, a limited number of studies have investigated sex differences in neonatal brain structure. This gap is surprising as the prenatal and neonatal periods are amongst the most rapid periods of brain development (17-19) and are likely critical windows for understanding sex differences in brain development. Moreover, given that brain development is highly dynamic during the first few weeks of life, existing findings from later stages of infancy cannot necessarily be extrapolated to the neonatal period. Neonatal research also provides a pivotal opportunity to understand the origins of sex differences in the brain and, specifically, the role of prenatal development in shaping these differences. To address this knowledge gap, we leveraged a sample of 514 healthy, term-born, singleton newborns (236 birth-assigned females, 278 birth-assigned males) from the developing Human Connectome Project (dHCP) to assess sex differences in neonatal global and regional brain volumes.

## Results

### Sample Characteristics

Welch’s two-sample t-tests showed no significant differences between males and females in gestational age at birth, postnatal age at scan, postconceptional age at scan, maternal age, or paternal age. There was a significant difference in birth weight and head circumference, both of which were greater in males (Table 1).

**Table 1.** Sample Characteristics.

|  | **Mean**<br>**(SD) All** | **Mean (SD)**<br>**Males** | **Mean (SD)**<br>**Females** | ***p*<sub>FDR</sub>** | **Cohen's *d*** |
| --- | --- | --- | --- | --- | --- |
| Gestational age at birth (weeks) | 40.15 (1.17) | 40.11 (1.14) | 40.21 (1.20) | 0.502 | 0.08 |
| Postnatal age at scan (days) | 8.69 (8.20) | 8.24 (7.85) | 9.21 (8.58) | 0.315 | 0.12 |
| Postconceptional age at scan (weeks) | 41.39 (1.62) | 41.29 (1.57) | 41.52 (1.67) | 0.233 | 0.15 |
| Birth weight (kg) | 3.44 (0.48) | 3.50 (0.46) | 3.36 (0.49) | 0.004 | 0.30 |
| Head circumference (cm) | 33.14<br>(1.64) | 35.49 (1.55) | 34.91 (1.69) | <0.001 | 0.36 |
| Maternal age (years) | 33.64<br>(4.81) | 33.62 (4.65) | 33.67 (4.99) | 0.910 | 0.01 |
| Paternal age (years) | 36.03<br>(6.15) | 35.82 (6.31) | 36.28 (5.96) | 0.502 | 0.07 |

*Note.* All *p*-values are FDR-corrected across the 7 analyses.

### Global Analyses

ANCOVA models were used to test for sex differences in global and regional brain volumes. After controlling for postconceptional age at scan and correcting for multiple comparisons, all global brain volumes (Table 2 and Figure 1) were larger in males than in females. All these differences remained significant after controlling for birth weight, except for the sex difference in cerebrospinal fluid (Table 3). After controlling for total brain volume in place of birth weight, males had larger total white matter volumes than females, whereas females had larger cortical gray matter volumes than males. There was no sex difference in total subcortical gray matter volumes (Table 3).

**Table 2.**
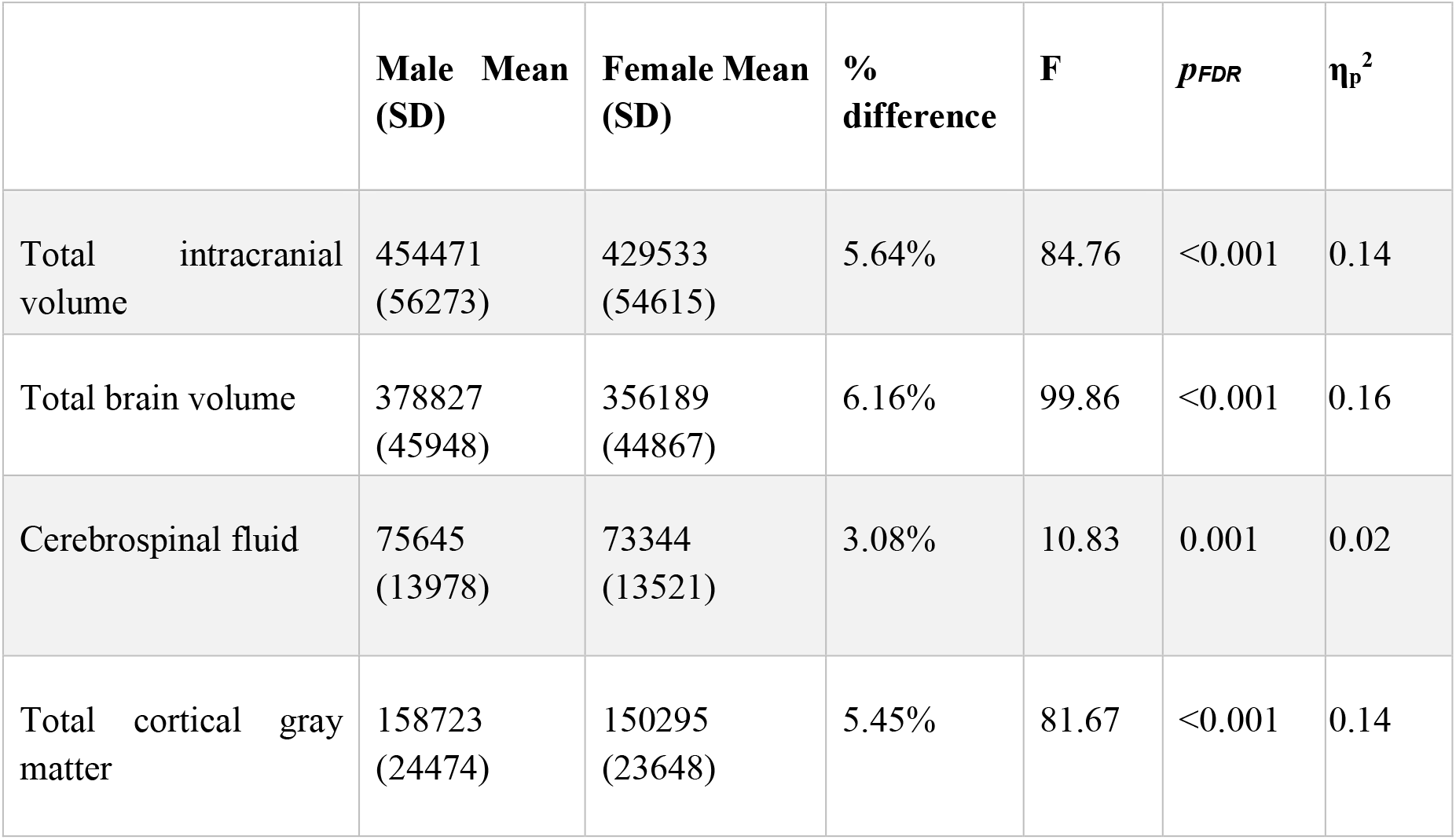
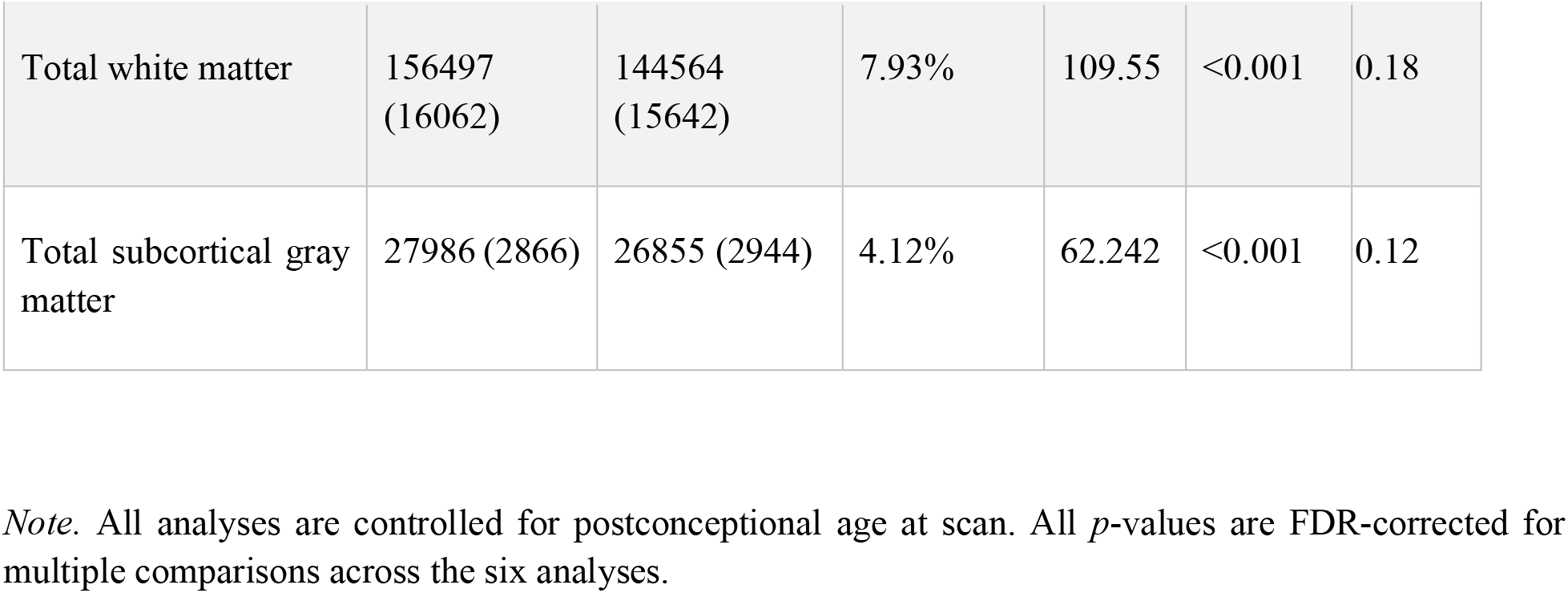
Absolute sex differences in global brain volumes (mm^3^)

|  | **Male Mean**<br>**(SD)** | **Female Mean**<br>**(SD)** | **%**<br>**difference** | **F** | ***p*<sub>FDR</sub>** | **$\eta_p^2$** |
| --- | --- | --- | --- | --- | --- | --- |
| Total intracranial volume | 454471<br>(56273) | 429533<br>(54615) | 5.64% | 84.76 | <0.001 | 0.14 |
| Total brain volume | 378827<br>(45948) | 356189<br>(44867) | 6.16% | 99.86 | <0.001 | 0.16 |
| Cerebrospinal fluid | 75645<br>(13978) | 73344<br>(13521) | 3.08% | 10.83 | 0.001 | 0.02 |
| Total cortical gray matter | 158723<br>(24474) | 150295<br>(23648) | 5.45% | 81.67 | <0.001 | 0.14 |
| Total white matter | 156497<br>(16062) | 144564<br>(15642) | 7.93% | 109.55 | <0.001 | 0.18 |
| Total subcortical gray matter | 27986 (2866) | 26855 (2944) | 4.12% | 62.242 | <0.001 | 0.12 |

*Note.* All analyses are controlled for postconceptional age at scan. All $p$ -values are FDR-corrected for multiple comparisons across the six analyses.

**Table 3.**
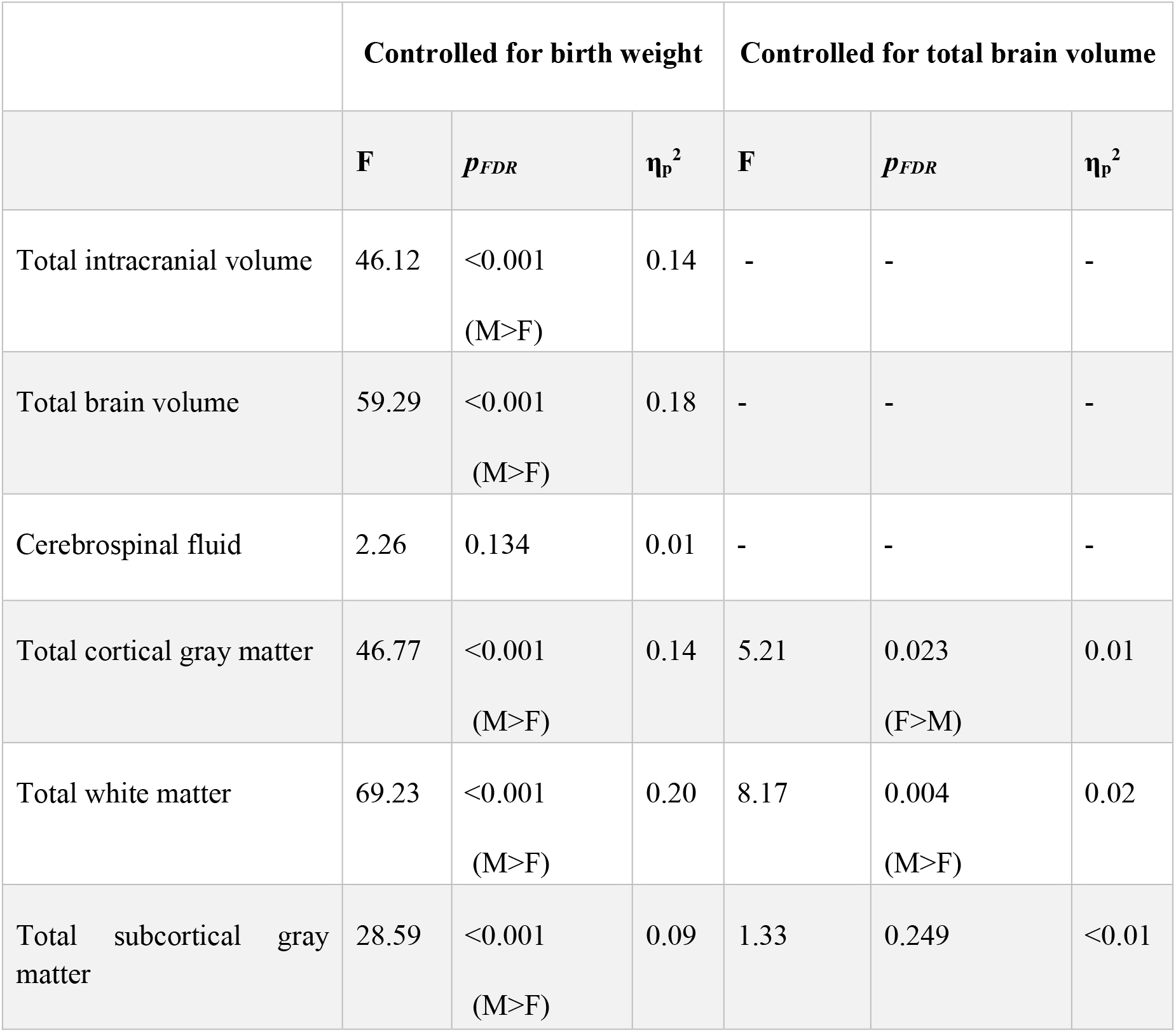
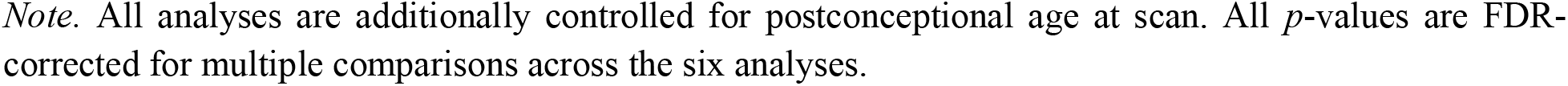
Sex differences in global brain volumes controlled for birth weight or total brain volume.

|  | Controlled for birth weight |  |  | Controlled for total brain volume |  |  |
| --- | --- | --- | --- | --- | --- | --- |
|  | **F** | **$p_{FDR}$** | **$\eta_p^2$** | **F** | **$p_{FDR}$** | **$\eta_p^2$** |
| Total intracranial volume | 46.12 | <0.001<br>(M>F) | 0.14 | - | - | - |
| Total brain volume | 59.29 | <0.001<br>(M>F) | 0.18 | - | - | - |
| Cerebrospinal fluid | 2.26 | 0.134 | 0.01 | - | - | - |
| Total cortical gray matter | 46.77 | <0.001<br>(M>F) | 0.14 | 5.21 | 0.023<br>(F>M) | 0.01 |
| Total white matter | 69.23 | <0.001<br>(M>F) | 0.20 | 8.17 | 0.004<br>(M>F) | 0.02 |
| Total subcortical gray matter | 28.59 | <0.001<br>(M>F) | 0.09 | 1.33 | 0.249 | <0.01 |

*Note.* All analyses are additionally controlled for postconceptional age at scan. All $p$ -values are FDR-corrected for multiple comparisons across the six analyses.

**Figure 1.**
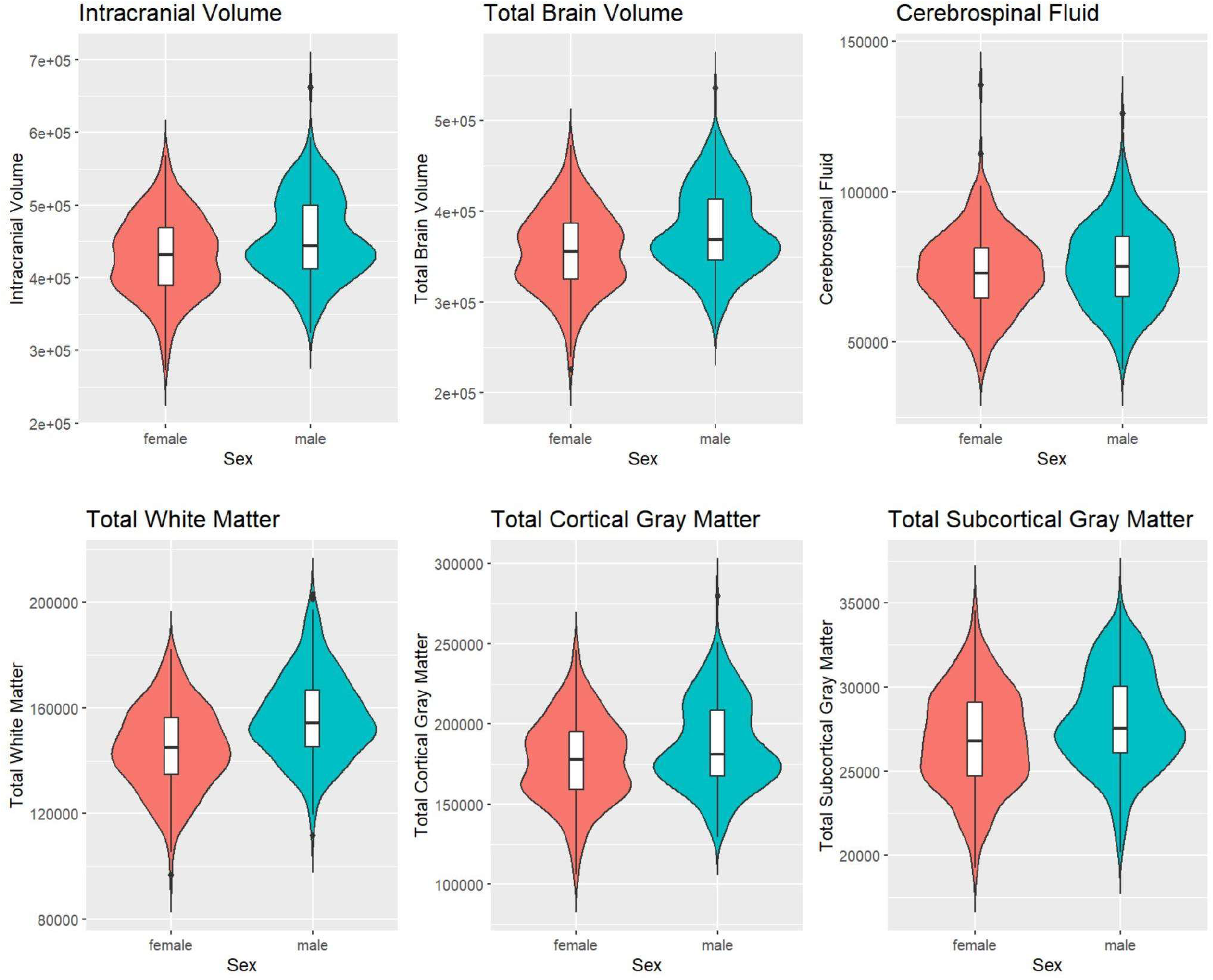
Absolute sex differences in global brain volumes

### Regional Analysis

After controlling for postconceptional age at scan, all regional volumes were larger in males (all FDR-corrected *p*<0.01). Full results of this analysis are reported in Supplementary Table S1. After additionally controlling for total brain volume, female>male sex differences were observed in 7 regions, including the white matter volume of the corpus callosum and the gray matter volumes of the bilateral parahippocampal gyri (posterior parts), left anterior cingulate gyrus, bilateral parietal lobes, and left caudate nucleus (Table 4). Male>female gray matter regions were observed in 2 regions, including the right medial and inferior temporal gyrus (posterior part) and right subthalamic nucleus (Table 5). These results are summarised in Tables 4 and 5 and visualised in Figure 2. Full results are reported in Supplementary Table S2.

**Table 4.** Significantly larger regions in females compared to males (female>male) after controlling for total brain volume.

| Region | Male Mean (SE) | Female Mean (SE) | F | $p_{FDR}$ | $\eta^2_p$ |
| --- | --- | --- | --- | --- | --- |
| Right parahippocampal gyrus (posterior part) | 815 (5.34) | 838 (5.84) | 7.81 | 0.010 | 0.02 |
| Left parahippocampal gyrus (posterior part) | 798 (5.78) | 821 (6.32) | 6.89 | 0.015 | 0.01 |
| Left anterior cingulate gyrus | 1314 (11.00) | 1352 (12.10) | 4.88 | 0.042 | 0.01 |
| Right Parietal Lobe | 18319 (50.10) | 18563 (54.80) | 9.95 | 0.003 | 0.020 |
| Left Parietal Lobe | 18454 (49.10) | 18685 (53.70) | 9.26 | 0.004 | 0.020 |
| Left Caudate Nucleus | 1879 (10.70) | 1921 (11.70) | 2.97 | 0.018 | 0.012 |
| Corpus Callosum | 2909 (18.90) | 2979 (20.70) | 5.82 | 0.026 | 0.011 |

*Note.* All $p$ -values are FDR-corrected for multiple comparisons across 48 regions. Means values are estimated marginal (EM) means controlled for total brain volume and postconceptional age at scan (absolute means are reported in supplementary materials).

**Table 5.**
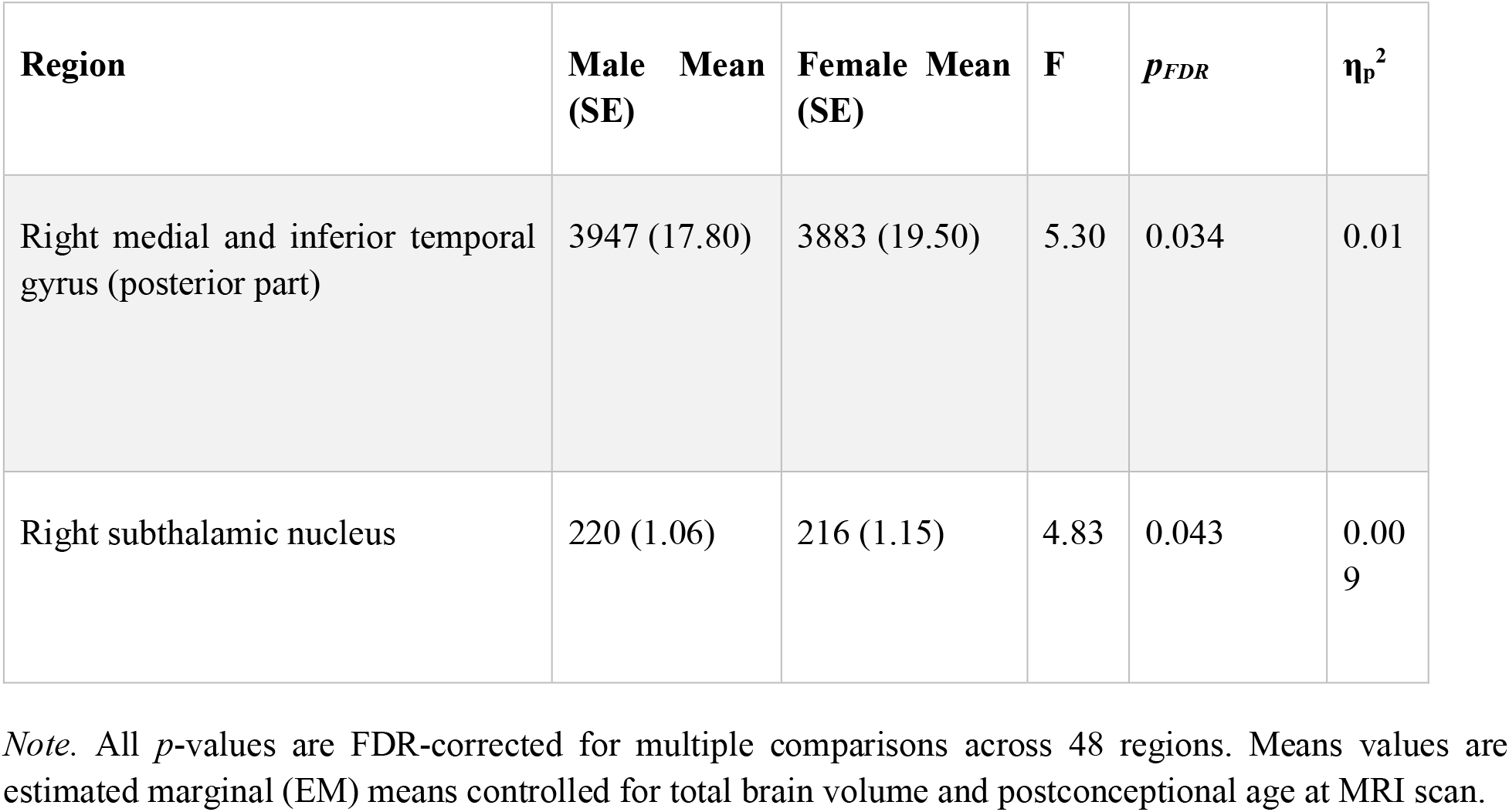
Significantly larger regions in males compared to females (male>female) after controlling for total brain volume.

| Region | Male Mean (SE) | Female Mean (SE) | F | $p_{FDR}$ | $\eta_p^2$ |
| --- | --- | --- | --- | --- | --- |
| Right medial and inferior temporal gyrus (posterior part) | 3947 (17.80) | 3883 (19.50) | 5.30 | 0.034 | 0.01 |
| Right subthalamic nucleus | 220 (1.06) | 216 (1.15) | 4.83 | 0.043 | 0.009 |

*Note.* All $p$ -values are FDR-corrected for multiple comparisons across 48 regions. Means values are estimated marginal (EM) means controlled for total brain volume and postconceptional age at MRI scan.

**Figure 2.**
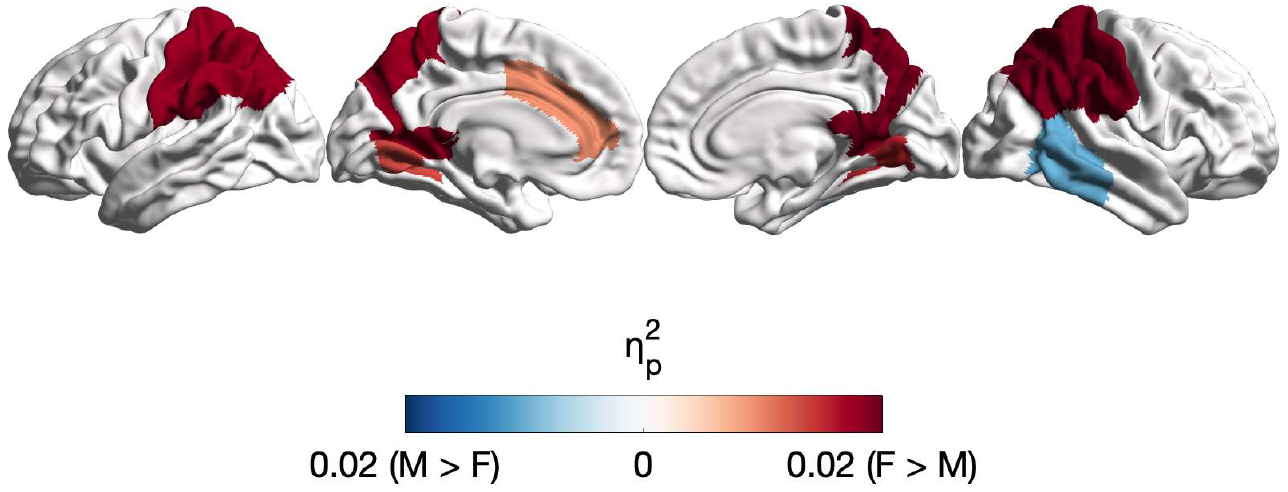
Regional sex differences by effect size after controlling for total brain volume *Note*. Figure 2 depicts **η_p_^2^** values of regions showing significant sex differences (*pFDR*<0.05) after controlling for total brain volume projected on a 32k Conte69 midthickness.

Full results for the model controlling for intracranial volume in place of total brain volume are reported in Supplementary Table S3 and Supplementary Figure S1. To summarise, female>male differences remained in the corpus callosum and left parahippocampal gyrus (posterior parts), and male>female differences remained in the medial and inferior temporal gyri and right subthalamic nucleus. A number of additional male>female differences were observed in the bilateral insula, bilateral amygdala, left subthalamic nucleus, bilateral superior temporal gyrus (middle part), left anterior temporal lobe (lateral part), and right frontal lobe.

## Discussion

In this research, we identified a number of global and regional on-average sex differences in neonatal brain volumes. All absolute global brain volumes were larger in males with large effect sizes, even after controlling for birth weight. After controlling for total brain volume, females showed increased total cortical gray matter volumes while males showed increased total white matter volumes. After controlling for total brain volume, various regional female>male (e.g., corpus callosum, posterior parts of the bilateral parahippocampal gyri, parietal lobes, left caudate nucleus, left anterior cingulate gyrus) and male>female (e.g., posterior right medial and inferior temporal gyrus) differences were identified with small effect sizes. As discussed further below, these findings suggest that several sex differences observed later in life are already present at birth.

First, we replicated the consistently reported finding that males have larger total brain (by 6.16%) and intracranial (by 5.64%) volumes than females, even after controlling for birth weight. The presence and magnitude of these differences is largely consistent with prior research in early infancy (11-14). These findings therefore confirm that sex differences in total brain volume are present from birth and are not fully accounted for by differences in body size. It is noteworthy that a meta-analysis (1) has previously reported 12% larger intracranial and 10.8% larger total brain volumes in males than females across the lifespan. Thus, although present at birth, these sex differences appear to increase in magnitude over the course of development. After controlling for total brain volume, females had larger total cortical gray matter volumes whilst males had larger total white matter volumes. This findings is largely consistent with research in later life stages (20-26). Collectively, these findings suggest that sex differences in global brain volumes are present from birth and are observed consistently throughout subsequent life stages.

After controlling for total brain volume, females had larger gray matter volumes in regions such as the bilateral parahippocampal gyri (posterior parts), left anterior cingulate gyrus, bilateral parietal lobes, and left caudate nucleus. Greater parietal lobe volumes in females have also been previously reported in early infancy (11, 16). Moreover, adult females show higher gray to white matter ratios (27-29) and greater cortical thickness in the parietal lobe (30-33) than adult males. Interestingly, prior work has also suggested a negative association between adolescent circulating testosterone levels and parietal lobe volumes (34).

A larger anterior cingulate cortex in females has previously been reported in early infancy (12, 15) and in a large sample of 2,328 adults (35). Similarly, a previous meta-analysis has reported larger volumes in the posterior parts of the parahippocampal gyrus in females (1). Numerous studies across the lifespan (34, 36-41), including research in young infants (14, 15) have also reported a larger caudate nucleus in females. The caudate nucleus, part of the basal ganglia, shows a high density of sex-steroid receptors (42, 43). Moreover, the caudate nucleus has been implicated in a number of conditions that show sex differences in their prevalence, such as ADHD (44, 45), Tourrete’s syndrome (46), depression (47, 48), and autism (49).

Finally, female newborns had relatively larger white matter volumes in the corpus callosum. An extensive body of previous research across various life stages supports the present findings (20, 50-54). It has been suggested that a larger corpus callosum may explain the lower hemispheric asymmetry observed in females (55, 56). Aspects of the corpus callosum, including its lateralisation and symmetry, also show associations with fetal testosterone levels (57). Importantly, the corpus callosum has been implicated in conditions that show sex differences and manifest during early childhood (58, 59).

On the other hand, males showed greater gray matter volumes in the subthalamic nucleus and the right medial and inferior temporal gyri (posterior parts) after controlling for total brain volume. Sex differences in the subthalamic nucleus have not been reported by prior research in later life, indicating that this sex difference might be unique to the neonatal stage. Regarding the medial and inferior temporal gyri, research in adolescents and adults corresponds with the present neonatal findings (26, 60). However, another study in early infancy has reported that the posterior parts of the medial and inferior temporal gyri were larger in females. It is important to note that the sample in this prior study included pre-term and twin infants, who are known to show different brain phenotypes and developmental trajectories compared to term-born, singleton infants (61, 62). These differences in sample characteristics, the use of a voxel-wise (tensor-based morphometry) rather than region-wise approach, and controlling for intracranial rather than total brain volume might explain the discrepancies with the present findings.

The decision to control for intracranial or total brain volume is an important one as the two approaches often yield different results, which perhaps stands as a leading source for inconsistency between existing studies in the field (2). In this study, we present findings from both approaches to facilitate cross-study comparability. Sex differences that were consistent across the two analyses included female>male differences in the corpus callosum and left parahippocampal gyrus (posterior parts), and male>female differences in the right subthalamic nucleus and right medial and inferior temporal gyrus (posterior parts). However, controlling for intracranial volume also yielded a number of additional male>female differences in regions such as the bilateral amygdala, bilateral insula, and right frontal lobe – all of which have also been documented by prior research (1, 35). This pattern aligns with prior studies wherein controlling for intracranial rather than total brain volume typically shows a greater number of male>female differences (63, 64). The trend likely links to our finding that males continue to have larger total brain volumes even after controlling for intracranial volume, which might explain why male>female differences attenuate when controlling for total brain volume itself.

More broadly, three patterns appear to emerge by synthesising the findings of the present neonatal research with those from later life stages: (a) some sex differences observed throughout the lifespan appear to be present from birth; (b) some sex differences are absent at birth but present in later development; and (c) some sex differences are present at birth but absent in later development. Pattern (a) appears to be most prevalent in our findings, having been observed in all global brain volumes as well as various regional volumes (e.g., caudate nucleus, anterior cingulate cortex, corpus callosum, etc.). It has previously been proposed that sex differences can be categorised as either “persistent”, such that they are established early in development and persist throughout the lifespan, or “transient”, such that they are temporary to a specific developmental period (65). Under this framework, the findings identified in pattern (a) can be classified as persistent sex differences, although these differences might still be dynamic over development. For instance, the sex difference in brain size is persistent in the sense that it is present from birth, but dynamic in the sense that it increases in magnitude over the course of development.

Regarding pattern (b), sex differences typically observed in adults that we did not observe in this neonatal sample are seen in regions such as the hippocampus and fusiform gyrus (1, 35). These sex differences might manifest as a result of both environmental influences as well as biological factors that unfold over development. Findings falling under pattern (c) include the subthalamic nucleus and can be understood as transient sex differences that might emerge as a result of short-term effects of prenatal processes. Although these differences are no longer observed during later development, they might play some initial role in instigating sex-specific developmental trajectories. Going forward, it will be important to verify these patterns via further longitudinal research on sex differences over the lifespan. Recent work on brain structural changes throughout the lifespan (17) and subcortical development during early childhood (14) set examples for future research to build upon.

There are important considerations that need to be taken into account when interpreting the findings of this study. First, scanning infants soon after birth minimises, but does not entirely eliminate, the influence of environmental factors. There remains the possibility that both prenatal (e.g., maternal drug/alcohol exposure) and postnatal (e.g., parental interaction with a newborn) environmental influences are present at this stage. However, whether and how these factors contribute to sex differences in neonatal brain development is not fully understood (66). Second, whilst it is reasonable to speculate that these sex differences may be influenced by prenatal factors (such as fetal testosterone), it is important to note that our findings do not establish any causal relationships between the two. Third, there might be a delayed effect of some prenatal biological processes, with their outcomes manifesting only gradually over development (7, 67). This suggests that neonatal research might capture only those effects that are immediately observable, potentially missing later-emerging effects. Fourth, sex differences in brain structure are not necessarily synonymous with sex differences in brain function or behaviour (41, 68). Further research directly examining these links will be essential to understanding whether the present findings have any implications for sex differences in behaviour and cognition. Fifth, given that definitions of regions can differ by atlas, cross-study compatibility of regional differences can be compromised (69). Finally, the present research examines only one of the many ways the brain can differ between males and females. Further research employing other neuroanatomical and functional measures will be critical to achieving a comprehensive insight into sex differences in the neonatal brain.

Strengths of the present research include the relatively large sample size. Importantly, the majority of infants were scanned within the first few days of birth, allowing us to capture the early neonatal period prior to extensive postnatal environmental influences. Moreover, the dHCP structural pre-processing pipeline (70) used in this research is optimised for the neonatal brain and overcomes several challenges typically encountered in neonatal brain imaging (e.g., partial volume effects, low tissue contrast, motion artefacts, etc.). The pipeline’s output also shows high correspondence with manual assessments of tissue boundaries.

In conclusion, sex differences are well-evidenced across later development, but remain significantly underexplored during the neonatal period. While postnatal development likely amplifies sex differences in the brain, our findings demonstrate that several global and regional differences are present from birth. The early emergence of these differences supports the hypothesis that prenatal factors play a pivotal role in shaping sex differences in the brain. Importantly, these early-emerging differences might influence neurobiological development and explain sex differences in the prevalence of brain-based neuropsychiatric conditions. Going forward, understanding this link should be an important research priority in order to help tailor early diagnostic and support strategies based on sex.

## Methods and Materials

### Participants

Participants were recruited as part of the developing Human Connectome Project (dHCP) (67). which was ethically approved by the UK National Research Ethics Authority (14/LO/1169). The dHCP contains data from 783 newborn infants (71). The exclusion criteria employed in this study included preterm births (<37 weeks gestational age), multiple births, the presence of brain anomalies in the scan with likely analytical and clinical significance (determined by an expert perinatal neuroradiologist), a postnatal age >28 days at the time of the scan, and pregnancy or neonatal clinical complications. The final sample used in this research consisted of 514 (236 birth-assigned females, 278 birth-assigned males) healthy, term-born, singleton infants scanned within the first 28 days of life (see Table 1 in Results for sample characteristics). Figure 3 shows the distribution of infant postnatal age at the time of the MRI scan, indicating that most infants were scanned within the first few days after birth.

**Figure 3.**
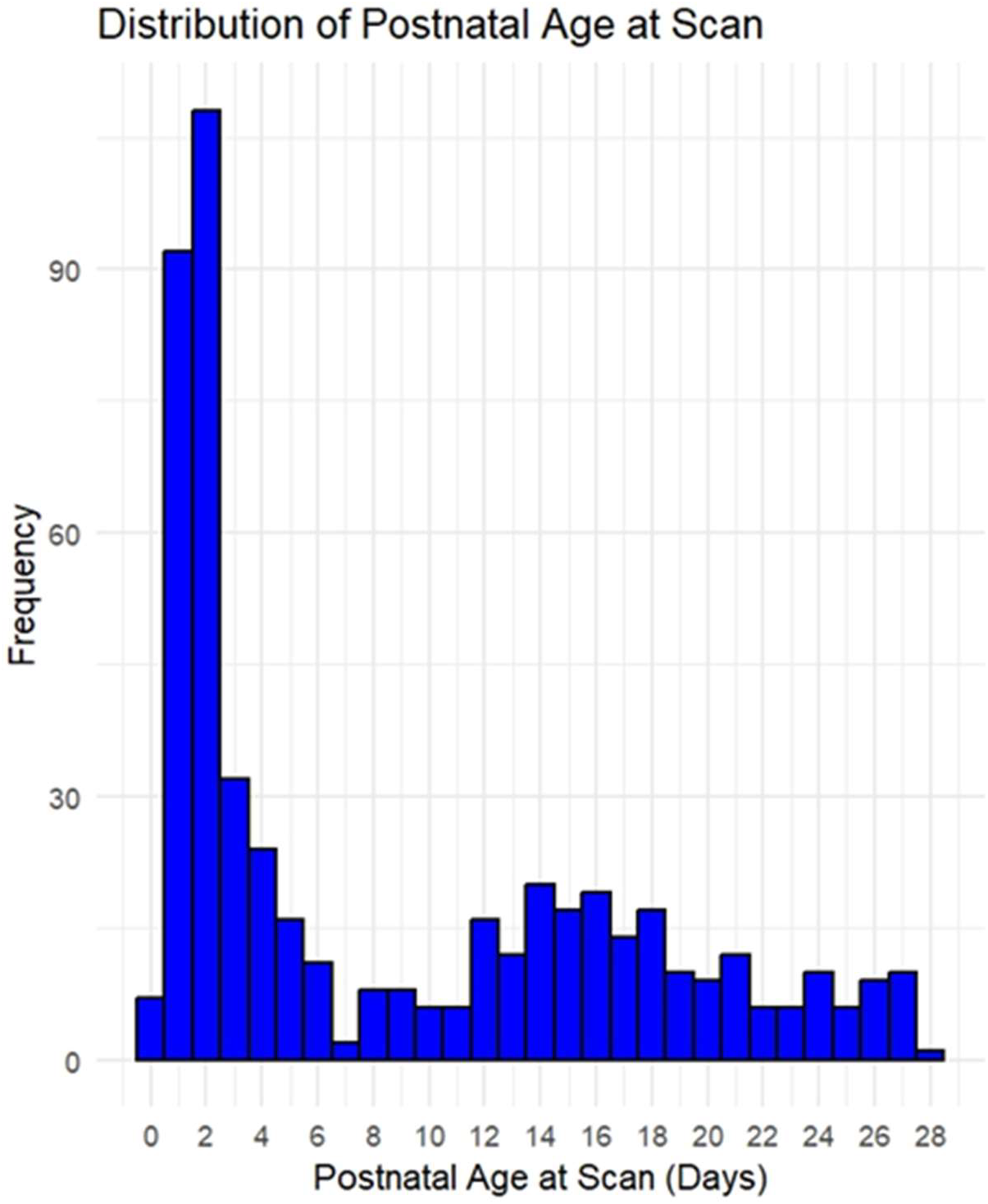
Distribution of Postnatal Age at Scan

### Data Acquisition

Data collection took place at the Evelina Newborn Imaging Centre, Evelina London Children’s Hospital. Data was acquired on a 3-Tesla Philips Achieva system (Philips Medical Systems) using the dHCP neonatal brain imaging system, which included a neonatal 32 channel phased array head coil and a customised patient handling system (Rapid Biomedical GmbH, Rimpar, Germany, 72). Infants were scanned without sedation after being fed and swaddled in a vacuum-evacuated blanket. For auditory protection, infants were equipped with earplugs (President Putty, Coltene Whaledent, Mahwah, NJ, USA) and neonatal earmuffs (MiniMuffs, Natus Medical Inc., San Carlos, CA, USA). Heart rate, oxygen saturation, and temperature were monitored throughout the scan by a paediatrician or neonatal nurse (71).

Anatomical data acquisition was conducted according to the specifications in the dHCP protocol (71). The imaging parameters were optimised to maximise contrast-to-noise ratio using a Cramer Rao Lower bound approach (Lankford and Does, 2013). Nominal relaxation times were set at T1/T2: 1800/150ms for gray matter and at T1/T2: 2500/250ms for white matter (73). T2-weighted and T1-weighted inversion recovery Fast Spin Echo (FSE) images were obtained in sagittal and axial planes. In-plane resolution was set at 0.8 × 0.8 mm^2^, with a slice thickness of 1.6 mm with 0.8 mm overlap. T1-weighted sagittal images used a slice overlap of 0.74mm. Other parameters were as follows - T2-weighted images: TR/TE = 12000/156 ms, SENSE factor 2.11 (axial) and 2.60 (sagittal); T1-weighted images: TR/TI/TE = 4795/1740/8.7 ms, SENSE factor 2.27 (axial) and 2.66 (sagittal). Additionally, 3D MPRAGE images were acquired using the following parameters: isotropic resolution = 0.8mm, TR/TI/TE = 11/1400/4.6 ms, SENSE factor 1.2 RL (Right-Left). These acquisitions were optimised for volumetric analysis using a motion correction algorithm, and transverse and sagittal images were fused into a single 3D volume for high resolution and accurate volume estimation (74).

### Data Preprocessing

The developing Human Connectome Project structural preprocessing pipeline was used for pre-processing the MRI data (70). To summarise, the T2-weighted image was first motion-corrected, bias-corrected and brain-extracted using the Brain Extraction tool (75). Next, a probabilistic tissue atlas was registered to the bias-corrected T2 image. Initial segmentation into different tissue types (i.e., cerebrospinal fluid, white matter, cortical gray matter, and subcortical gray matter) was performed using the Draw-EM algorithm (76). Labelled atlases (77) were then registered to the subject’s images via a multi-channel registration process, using both GM probability maps from the initial segmentation and intensity T2-weighted images. The resulting segmentation consisted of 87 gray and white matter structures (see 76-78).

### Data Analysis

Statistical analysis were conducted on R (version 4.3.3, 2024-02-29), using the packages rstatix, tidyverse, effectsize, and ggplot2. Analysis of Covariance (ANCOVA) models were used to test for sex differences in brain volumes. Postconceptional age at the time of the MRI scan was used as a covariate in all models. To account for size, birth weight was included as a covariate in global analyses, while total brain volume was included as a covariate in regional and total gray/white matter analyses (apart from when testing for absolute sex differences). The measure of total brain volume was derived by summing the volumes of all cortical and subcortical structures excluding the ventricles and cerebrospinal fluid. To facilitate cross-study comparability with studies that have used intracranial volume as a covariate, further regional analyses were conducted controlling for intracranial rather than total brain volume. The measure of total intracranial volume was derived by adding cerebrospinal fluid volume to total brain volume. Regional analyses focused primarily on gray matter volumes and were conducted on 47 cortical and subcortical gray matter regions (73). The white matter volume of the corpus callosum, however, was also included in the analysis since sex differences in corpus callosum volume are of key interest due to its critical role in inter-hemispheric connectivity (51, 79). Analyses were corrected for multiple comparisons using the Benjamini-Hochberg FDR correction (80) with a significance threshold of 0.05. FDR corrections were run separately for global volumes (6 tests) and regional volumes (48 tests) for each ANCOVA model. Additionally, Welch’s two-sample t-tests were used to assess sex differences in sample characteristics.

## Acknowledgements

We would like to thank the developing Human Connectome Project consortium for making available the data used in this analysis. We would also like to thank Michael V. Lombardo for providing constructive feedback on the draft manuscript of this paper, and Saashi A. Bedford for assisting with organising the processing of the data.

## Author contributions

Design and conceptualisation: Y.T.K., R.A.I.B., A.T., S.B.-C., C.A. Data processing: A.T., Y.T.K., L.D. Analysis: Y.T.K. Visualisation: M.A.R. Supervision: S.B.-C., C.A. Writing - original draft: Y.T.K. Writing - reviewing and editing: All authors.

## Data availability

All data needed to evaluate the conclusions in the paper are present in the paper and/or the Supplementary Materials.

## APEX Consortium

Deep Adhya, Carrie Allison, Bonnie Ayeung, Rosie Bamford, Simon Baron-Cohen, Richard A. I. Bethlehem, Tal Biron-Shental, Graham Burton, Wendy Cowell, Jonathan Davies, Dorothea L. Floris, Alice Franklin, Lidia Gabis, Daniel Geschwind, David M. Greenberg, Yuanjun Gu, Alexandra Havdahl, Alexander Heazell, Rosemary J. Holt, Matthew Hurles, Yumnah T. Khan, Meng-Chuan Lai, Madeline Lancaster, Michael V. Lombardo, Hilary Martin, Jose Gonzalez Martinez, Jonathan Mill, Mahmoud Musa, Kathy Niakan, Adam Pavlinek, Lucia Dutan Polit, Marcin A. Radecki, David Rowitch, Jenifer Sakai, Laura Sichlinger, Deepak Srivastava, Alex Tsompanidis, Florina Uzefovsky, Varun Warrier, Elizabeth M. Weir, Xinhe Zhang.

