## Supplementary Materials for "Sex Differences in Human Brain Structure at Birth" (from another version of this preprint: version 1, 2024-06-20)

**Supplementary Materials: Sex Differences in Human Brain Structure at Birth**

**Supplementary Table S1.** *Absolute Sex Differences in Regional Volumes*

| **Region** | **Male Mean (SD)** | **Female Mean (SD)** | **F** | $p_{FDR}$ | $\eta_p^2$ |
|---|---|---|---|---|---|
| Hippocampus left | 754.61 (108.54) | 710.08 (96.14) | 43.14 | <0.001 | 0.048 |
| Hippocampus right | 843.27 (116.79) | 794.69 (100.65) | 47.16 | <0.001 | 0.052 |
| Amygdala left | 493.06 (57.28) | 464.99 (53.31) | 57.96 | <0.001 | 0.066 |
| Amygdala right | 560.85 (69.89) | 528.73 (65.32) | 53.87 | <0.001 | 0.059 |
| Anterior temporal lobe medial part left | 950.06 (170.31) | 892.01 (155.54) | 37.71 | <0.001 | 0.035 |
| Anterior temporal lobe medial part right | 987.10 (179.93) | 931.49 (168.79) | 36.53535 | <0.001 | 0.030 |
| Anterior temporal lobe lateral part left | 1018.61 (195.79) | 949.41 (194.67) | 46.89 | <0.001 | 0.040 |
| Anterior temporal lobe lateral part right | 1038.42 (188.05) | 992.35 (197.97) | 27.64 | <0.001 | 0.021 |
| Parahippocampal gyrus left (anterior part) | 1152.14 (151.43) | 1092.46 (140.68) | 32.00 | <0.001 | 0.041 |

| | | | | | |
|---|---|---|---|---|---|
| Parahippocampal gyrus right (anterior part) | 1035.84 (154.27) | 992.35 (139.68 ) | 21.89 | <0.001 | 0.023 |
| Superior temporal gyrus left (middle part) | 2995.34 (485.12) | 2818.21 (499.37) | 62.24 | <0.001 | 0.044 |
| Superior temporal gyrus right (middle part) | 3060.42 (487.87) | 2882.87 (498.09) | 60.88 | <0.001 | 0.044 |
| Medial and inferior temporal gyri left (anterior part) | 2513.80 (490.85) | 2364.28 (481.85) | 42.91 | <0.001 | 0.031 |
| Medial and inferior temporal gyri right (anterior part) | 2717.62 (536.70) | 2582.42 (505.43) | 38.76 | <0.001 | 0.024 |
| Lateral occipitotemporal gyrus left (anterior part) | 738.82 (136.94) | 703.06 (147.25 ) | 21.75 | <0.001 | 0.020 |
| Lateral occipitotemporal gyrus right (anterior part) | 782.66 (161.29) | 742.26 (161.44 ) | 23.81 | <0.001 | 0.021 |
| Cerebellum left | 13274.96 (2187.81) | 12914.43 (2163.71) | 35.08 | <0.001 | 0.012 |
| Cerebellum right | 12983.41 (2142.42) | 12573.07 (2056.700) | 41.06 | <0.001 | 0.016 |
| Brainstem | 6711.41 (631.78) | 6489.43 (631.78) | 47.65 | <0.001 | 0.038 |

| | | | | | |
|---|---|---|---|---|---|
| Insula right | 1768.60 (274.71) | 1649.47 (258.102) | 76.54 | <0.001 | 0.058 |
| Insula left | 1956.75 (291.77) | 1822.12 (261.383) | 78.74 | <0.001 | 0.067 |
| Occipital lobe right | 12211.90 (2041.58 ) | 11665.80 (1940.91) | 40.74 | <0.001 | 0.027 |
| Occipital lobe left | 12332.42 (2036.91) | 11787.81 (2015.79) | 43.93 | <0.001 | 0.027 |
| Parahippocampal gyri right (posterior part) | 837.90 (144.29) | 811.61 (132.87 ) | 15.76 | <0.001 | 0.012 |
| Parahippocampal gyri rleft (posterior part) | 816.78 (127.02) | 798.84 (130.97 ) | 8.41 | 0.004 | 0.007 |
| Latetal occipitotemporal gyus right (posterior part) | 1027.06 (209.30) | 988.73 (207.87 ) | 19.02 | <0.001 | 0.013 |
| Lateral occipitotemporal gyrus left (posterior part) | 978.36 (201.88) | 937.59 (194.25) | 16.69 | <0.001 | 0.014 |
| Medial and inferior temporal gyri right (posterior part) | 4058.45 (713.17) | 3751.90 (696.98) | 88.78 | <0.001 | 0.061 |
| Medial and inferior temporal gyri left (posterior part) | 4159.01 (753.13) | 3864.00 (711.37) | 76.32 | <0.001 | 0.052 |

| | | | | | |
|---|---|---|---|---|---|
| Superior temporal gyrus right (posterior part) | 1295.30 (256.28) | 1237.55 (233.245) | 25.52 | <0.001 | 0.019 |
| Superior temporal gyrus left (posterior part) | 1144.14 (239.22) | 1094.56 (211.38) | 23.94 | <0.001 | 0.017 |
| Cingulate gyrus right (anterior part) | 1544.77 (252.26) | 1433.81 (247.70) | 44.87 | <0.001 | 0.051 |
| Cingulate gyrus left (anterior part) | 1364.15 (234.38) | 1293.74 (237.47) | 17.41 | <0.001 | 0.023 |
| Cingulate gyrus right (posterior part) | 1787.37 (309.72) | 1652.96 (295.31) | 52.99 | <0.001 | 0.053 |
| Cingulate gyrus left (posterior part) | 1748.06 (301.37) | 1643.19 (299.28) | 40.26 | <0.001 | 0.036 |
| Frontal lobe right | 26574.61 (4274.314) | 24900.72 (4063.05) | 90.83 | <0.001 | 0.053 |
| Frontal lobe left | 26410.15 (4264.03) | 24880.23 (4104.48) | 80.80 | <0.001 | 0.044 |
| Parietal lobe right | 18784.53 (3027.97) | 18014.77 (2946.75) | 53.25 | <0.001 | 0.025 |
| Parietal lobe left | 18931.88 (3071.98) | 18122.12 (2947.58) | 55.85 | <0.001 | 0.027 |
| Caudate nucleus right | 1989.20 (261.90) | 1902.69 (265.61) | 27.21 | <0.001 | 0.028 |

| | | | | | |
|---|---|---|---|---|---|
| Caudate nucleus left | 1933.80 (256.78) | 1856.97 (275.68) | 22.02 | <0.001 | 0.022 |
| Thalamus right | 4245.96 (442.83) | 4102.72 (422.09) | 46.35 | <0.001 | 0.034 |
| Thalamus left | 4163.66 (433.66) | 4012.22 (423.85) | 53.84 | <0.001 | 0.039 |
| Subthalamic nucleus right | 224.42 (26.22) | 210.93 (26.55) | 66.62 | <0.001 | 0.072 |
| Subthalamic nucleus left | 187.45 (20.52) | 176.62 (23.76) | 48.81 | <0.001 | 0.065 |
| Lentiform nucleus right | 3223.46 (437.95) | 3074.19 (435.16) | 52.17 | <0.001 | 0.037 |
| Lentiform nucleus left | 3423.82 (421.25) | 3256.58 (416.04) | 60.40 | <0.001 | 0.048 |
| Corpus Callosum | 2994.48 (477.23) | 2878.41 (459.07) | 19.29 | <0.001 | 0.018 |

*Note.* All analyses are controlled for postconceptional age at scan. All *p*-values are FDR-corrected for multiple comparisons.

**Supplementary Table S2.** *Sex Differences in regional volumes after controlling for total brain volume*

| **Region** | **Male EM Mean (SE)** | **Female EM Mean (SE)** | **F** | $p_{FDR}$ | $\eta_p^2$ |
|---|---|---|---|---|---|
| Hippocampus left | 737 (4.52) | 731 (4.94) | 0.54 | 0.504 | 0.001 |

| | | | | | |
|---|---|---|---|---|---|
| Hippocampus right | 824 (4.66) | 817 (5.09) | 0.82 | 0.418 | 0.002 |
| Amygdala left | 483 (2.36) | 477 (2.58) | 2.91 | 0.119 | 0.006 |
| Amygdala right | 549 (2.89) | 542 (3.16) | 2.41 | 0.163 | 0.005 |
| Anterior temporal lobe medial part left | 927 (7.00) | 919 (7.66) | 0.67 | 0.467 | 0.001 |
| Anterior temporal lobe medial part right | 959 (6.39) | 965 (6.99) | 0.32 | 0.604 | 0.001 |
| Anterior temporal lobe lateral part left | 978 (8.26) | 994 (7.55) | 1.88 | 0.216 | 0.004 |
| Anterior temporal lobe lateral part right | 1013 (7.09) | 1022 (7.75) | 0.58 | 0.493 | 0.001 |
| Parahippocampal gyrus left (anterior part) | 1124 (6.95) | 1125 (7.60) | 0.01 | 0.493 | 0.000 |
| Parahippocampal gyrus right (anterior part) | 1008 (6.40) | 1025 (7.00) | 3.05 | 0.113 | 0.006 |
| Superior temporal gyrus left (middle part) | 2932 (15.60) | 2893 (17.10) | 2.60 | 0.145 | 0.005 |

| | | | | | |
|---|---|---|---|---|---|
| Superior temporal gyrus right (middle part) | 2991 (15.2) | 2964 (16.60) | 1.35 | 0.298 | 0.003 |
| Medial and inferior temporal gyri left (anterior part) | 2438 (15.20) | 2454 (16.70) | 0.47 | 0.530 | 0.001 |
| Medial and inferior temporal gyri right (anterior part) | 2641 (15.00) | 2673 (16.40) | 1.87 | 0.216 | 0.004 |
| Lateral occipitotemporal gyrus left (anterior part) | 718 (5.98) | 728 (6.54) | 1.21 | 0.322 | 0.002 |
| Lateral occipitotemporal gyrus right (anterior part) | 760 (6.56) | 769 (7.18) | 0.78 | 0.432 | 0.002 |
| Cerebellum left | 13061 (54.10) | 13166 (59.20) | 1.55 | 0.265 | 0.003 |
| Cerebellum right | 12782 (54.70) | 12811 (59.80) | 0.12 | 0.754 | 0.000 |
| Brainstem | 6593 (18.40) | 6629 (20.10) | 1.55 | 0.263 | 0.003 |

| | | | | | |
|---|---|---|---|---|---|
| Insula right | 1726 (8.29) | 1699 (9.06) | 4.35 | 0.056 | 0.008 |
| Insula left | 1908 (8.79) | 1880 (9.61) | 4.32 | 0.056 | 0.008 |
| Occipital lobe right | 11891 (54.30) | 12043 (59.40) | 3.27 | 0.102 | 0.006 |
| Occipital lobe left | 12014 (50.30) | 12163 (55.00) | 3.65 | 0.081 | 0.007 |
| Parahippocampal gyri right (posterior part) | 815 (5.34) | 838 (5.84) | 7.81 | 0.010 | 0.020 |
| Parahippocampal gyri left (posterior part) | 798 (5.78) | 821 (6.32) | 6.89 | 0.015 | 0.015 |
| Lateral occipitotemporal gyrus right (posterior part) | 1004 (8.12) | 1016 (8.88) | 1.07 | 0.350 | 0.002 |
| Lateral occipitotemporal gyrus left (posterior part) | 9.53 (8.65) | 967 (9.46) | 1.10 | 0.347 | 0.002 |

| | | | | | |
|---|---|---|---|---|---|
| Medial and inferior temporal gyri right (posterior part) | 3947 (17.80) | 3883 (19.50) | 5.30 | 0.034 | 0.010 |
| Medial and inferior temporal gyri left (posterior part) | 4044 (19.40) | 4000 (21.20) | 2.20 | 0.183 | 0.004 |
| Superior temporal gyrus right (posterior part) | 1262 (9.21) | 1277 (10.07) | 1.06 | 0.350 | 0.002 |
| Superior temporal gyrus left (posterior part) | 1118 (8.75) | 1126 (9.56) | 0.38 | 0.575 | 0.001 |
| Cingulate gyrus right (anterior part) | 1493 (10.00) | 1494 (11.00) | 0.006 | 0.948 | 0.000 |
| Cingulate gyrus left (anterior part) | 1314 (11.00) | 1352 (12.10) | 4.88 | 0.043 | 0.009 |
| Cingulate gyrus right (posterior part) | 1722 (10.00) | 1730 (10.90) | 0.27 | 0.636 | 0.001 |
| Cingulate gyrus left (posterior part) | 1688 (9.65) | 1714 (10.55) | 2.87 | 0.123 | 0.005 |
| Frontal lobe right | 25836 (60.90) | 25771 (66.59) | 0.48 | 0.530 | 0.001 |

| | | | | | |
|---|---|---|---|---|---|
| Frontal lobe left | 25679 (59.33) | 25742 (64.88) | 0.47 | 0.530 | 0.001 |
| Parietal lobe right | 18319 (50.10) | 18563 (54.80) | 9.95 | 0.003 | 0.020 |
| Parietal lobe left | 18454 (49.10) | 18685 (53.70) | 9.26 | 0.004 | 0.020 |
| Caudate nucleus right | 1936 (10.70) | 1965 (11.70) | 2.97 | 0.118 | 0.006 |
| Caudate nucleus left | 1879 (10.70) | 1921 (11.70) | 6.58 | 0.018 | 0.012 |
| Thalamus right | 4167 (12.10) | 4195 (13.30) | 2.18 | 0.184 | 0.004 |
| Thalamus left | 4087 (11.60) | 4102 (12.70) | 0.650 | 0.465 | 0.001 |
| Subthalamic nucleus right | 220 (1.06) | 216 (1.15) | 4.83 | 0.043 | 0.009 |
| Subthalamic nucleus left | 184 (1.05) | 181 (1.15) | 3.10 | 0.111 | 0.006 |
| Lentiform nucleus right | 3159 (13.90) | 3159 (13.90) | 0.19 | 0.695 | 0.009 |
| Lentiform nucleus left | 3358 (13.90) | 3334 (15.20) | 1.33 | 0.299 | 0.003 |

| | | | | | |
|---|---|---|---|---|---|
| Corpus Callosum | 2909 (18.90) | 2979 (20.70) | 5.82 | 0.026 | 0.011 |

*Note.* All analyses are controlled for postconceptional age at scan and total brain volume. All *p* values are FDR-corrected for multiple comparisons. Means values are estimated marginalised means corrected for total brain volume and postconceptional age at birth.

**Table S3.** *Sex Differences in regional volumes after controlling for intracranial volume*

| **Region** | **Male EM Mean (SE)** | **Female EM Mean (SE)** | **F** | $p_{FDR}$ | $\eta_p^2$ |
|---|---|---|---|---|---|
| Hippocampus left | 739 (4.56) | 729 (4.98) | 2.09 | 0.198 | 0.004 |
| Hippocampus right | 826 (4.71) | 814 (5.15) | 2.74 | 0.137 | 0.005 |
| Amygdala left | 484 (2.61) | 475 (2.61) | 5.99 | 0.023 | 0.010 |
| Amygdala right | 551 (2.89) | 541 (3.16) | 4.81 | 0.043 | 0.009 |
| Anterior temporal lobe medial part left | 931 (7.12) | 914 (7.78) | 2.52 | 0.156 | 0.001 |
| Anterior temporal lobe medial part right | 963 (6.46) | 960 (7.06) | 0.05 | 0.870 | 0.001 |
| Anterior temporal lobe lateral part left | 999 (7.70) | 973 (8.41) | 4.710 | 0.045 | 0.004 |
| Anterior temporal lobe lateral part right | 1017 (7.17) | 1017 (7.84) | 0.00 | 0.988 | 0.001 |

| | | | | | |
|---|---|---|---|---|---|
| Parahippocampal gyrus left (anterior part) | 1128 (7.00) | 1121 (7.65) | 0.33 | 0.638 | 0.000 |
| Parahippocampal gyrus right (anterior part) | 1012 (6.51) | 1021 (7.12) | 0.81 | 0.438 | 0.002 |
| Superior temporal gyrus left (middle part) | 2941 (15.86) | 2882 (17.32) | 5.94 | 0.024 | 0.010 |
| Superior temporal gyrus right (middle part) | 3005 (15.92) | 2949 (17.39) | 5.18 | 0.036 | 0.010 |
| Medial and inferior temporal gyri left (anterior part) | 2451 (16.05) | 2438 (17.54) | 0.30 | 0.654 | 0.001 |
| Medial and inferior temporal gyri right (anterior part) | 2653 (15.55) | 2659 (16.98) | 0.07 | 0.844 | 0.000 |
| Lateral occipitotemporal gyrus left (anterior part) | 722 (6.14) | 723 (6.71) | 0.03 | 0.902 | 0.000 |

| | | | | | |
|---|---|---|---|---|---|
| Lateral occipitotemporal gyrus right (anterior part) | 764.38 (6.75) | 763.79 (7.37) | <0.01 | 0.961 | 0.000 |
| Cerebellum left | 13092 (54.38) | 13131 (59.40) | 0.22 | 0.702 | 0.000 |
| Cerebellum right | 12812 (55.03) | 12776 (60.12) | 0.18 | 0.721 | 0.000 |
| Brainstem | 6608 (18.96) | 6611 (20.71) | 0.01 | 0.942 | 0.000 |
| Insula right | 1732 (8.50) | 1692 (9.29) | 9.25 | 0.004 | 0.020 |
| Insula left | 1916 (9.23) | 1871 (10.08) | 10.13 | 0.003 | 0.020 |
| Occipital lobe right | 11930 (54.94) | 11999 (60.02) | 0.666 | 0.482 | 0.001 |
| Occipital lobe left | 12051 (50.76) | 12120 (55.45) | 0.78 | 0.447 | 0.002 |
| Parahippocampal gyri right (posterior part) | 818 (5.39) | 835 (5.88) | 4.21 | 0.059 | 0.008 |

| | | | | | |
|---|---|---|---|---|---|
| Parahippocampal gyri left (posterior part) | 800 (5.75) | 819 (6.28) | 4.83 | 0.043 | 0.009 |
| Lateral occipitotemporal gyrus right (posterior part) | 1006 (8.06) | 1014 (8.80) | 0.38 | 0.612 | 0.001 |
| Lateral occipitotemporal gyrus left (posterior part) | 956 (8.58) | 964 (9.37) | 0.44 | 0.581 | 0.001 |
| Medial and inferior temporal gyri right (posterior part) | 3960 (17.94) | 3868 (19.60) | 10.94 | 0.002 | 0.010 |
| Medial and inferior temporal gyri left (posterior part) | 4060 (20.08) | 3980 (21.93) | 6.66 | 0.017 | 0.010 |
| Superior temporal gyrus right (posterior part) | 1267 (9.37) | 1270 (10.24) | 0.04 | 0.878 | 0.000 |
| Superior temporal gyrus left (posterior part) | 1122 (8.87) | 1120 (9.69) | 0.01 | 0.929 | 0.000 |

| | | | | | |
|---|---|---|---|---|---|
| Cingulate gyrus right (anterior part) | 1502 (10.60) | 1484 (11.58) | 1.29 | 0.315 | 0.003 |
| Cingulate gyrus left (anterior part) | 1323 (11.49) | 1342 (12.55) | 1.30 | 0.315 | 0.002 |
| Cingulate gyrus right (posterior part) | 1719 (11.58) | 1731 (10.60) | 0.61 | 0.503 | 0.001 |
| Cingulate gyrus left (posterior part) | 1731 (10.60) | 1704 (10.97) | 0.24 | 0.692 | 0.000 |
| Frontal lobe right | 25928 (66.82) | 25662 (72.99) | 6.70 | 0.016 | 0.010 |
| Frontal lobe left | 25768 (64.52) | 25637 (70.48) | 1.76 | 0.240 | 0.003 |
| Parietal lobe right | 18386 (55.12) | 18484 (60.22) | 1.33 | 0.314 | 0.003 |
| Parietal lobe left | 18517 (52.91) | 18611 (57.80) | 1.32 | 0.315 | 0.003 |
| Caudate nucleus right | 1946 (11.34) | 1954 (12.39) | 0.18 | 0.721 | 0.009 |
| Caudate nucleus left | 1889 (11.35) | 1909 (12.40) | 1.29 | 0.315 | 0.000 |
| Thalamus right | 4179 (12.77) | 4182 (13.96) | 0.02 | 0.906 | 0.003 |
| Thalamus left | 4098 (12.15) | 4090 (13.28) | 0.20 | 0.709 | 0.000 |
| Subthalamic nucleus right | 221 (1.08) | 215 (1.18) | 9.08 | 0.005 | 0.020 |

| | | | | | |
|---|---|---|---|---|---|
| Subthalamic nucleus left | 184 (1.07) | 180 (1.16) | 6.29 | 0.020 | 0.010 |
| Lentiform nucleus right | 3168 (14.10) | 3140 (15.40) | 1.72 | 0.241 | 0.003 |
| Lentiform nucleus left | 3368 (14.25) | 3322 (15.57) | 4.39 | 0.054 | 0.009 |
| Corpus Callosum | 2909 (17.59) | 2980 (19.21) | 6.90 | 0.015 | 0.010 |

*Note.* All analyses are controlled for postconceptional age and intracranial volume. All *p* values are FDR-corrected for multiple comparisons. Means values are estimated marginalised means corrected for intracranial volume and postconceptional age at birth.

**Supplementary Figure S1.**

*Regional sex differences by effect size after controlling for intracranial volume*

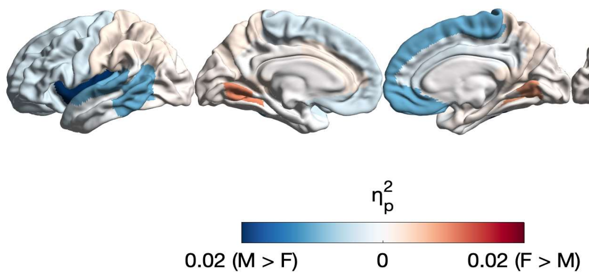

*Note.* Supplementary Figure 1 depicts $\boldsymbol{\eta_p}^2$ values of regions showing statistically significant sex differences (FDR-corrected $p < 0.05$) after controlling for intracranial volume.
